# Impact of sterilization on a conjugated polymer based bioelectronic patch

**DOI:** 10.1101/2021.01.19.427349

**Authors:** Yihan Yan, Lorenzo Travaglini, Kieran Lau, Jelena Rnjak-Kovacina, Minoo Eslami, Shaohua Yang, Antonio Lauto, David Officer, Damia Mawad

## Abstract

Translation into the clinic of organic bioelectronic devices having conjugated polymers as the active material will hinge on their long-term operation *in vivo*. This will require the device to be subject to clinically approved sterilization techniques without a deterioration in its physical and electronic properties. To date, there remains a gap in the literature addressing the impact of this critical pre-operative procedure on the properties of conjugated polymers. This study aims to address this gap by assessing the physical and electronic properties of a sterilized porous bioelectronic patch having polyaniline as the conjugated polymer. The patch was sterilized by autoclave, ethylene oxide and gamma (γ-) irradiation at 15, 25, and 50 kGy doses. Autoclaving resulted in cracking and macroscopic degradation of the patch, while patches sterilized by γ-irradiation at 50 kGy exhibited reduced mechanical and electronic properties, attributed to chain scission and non-uniform crosslinking caused by the high dose irradiation. Ethylene oxide and γ-irradiation at 15 and 25 kGy sterilization appeared to be the most effective at maintaining the mechanical and electronic properties of the patch, as well as inducing a minimal immune response as revealed by a receding fibrotic capsule after 4 weeks implantation. Our findings pave the way towards closing the gap for the translation of organic bioelectronic devices from acute to long-term *in vivo* models.

## 1. INTRODUCTION

Flexible organic bioelectronic devices with a conjugated polymer as the active material have been shown to effectively interface with biological tissues, demonstrated by their capability to stimulate or record electro-physiological signals.^1,2^ Their beneficial communication with the underlying organ is enabled by the conformity of the device and its intimate contact to the tissue.^3,4^ Significant developments have been achieved since the discovery of conjugated polymers, with the field shifting from *in vitro* substrates^5,6^ to conductive coatings of implantable electrodes,^7,8^ and more recently as active materials in advanced bioelectronic circuitry.^9–12^ Their functionality has been largely demonstrated in acute *in vivo* models, with promising results indicating that these organic bioelectronics are poised to have an impact in future medical applications.^3,13^ However, their translation into the clinic will require their sterilization, a critical preoperative process, without degrading their physical and electronic properties. Despite the tremendous amount of research conducted on organic bioelectronics, there remains a gap in the literature dedicated towards understanding the compatibility of conjugated materials with clinically approved sterilization techniques.^4^ Here, we investigate the effect of autoclave, ethylene oxide (EO), and gamma (γ) irradiation on the physical and electronic properties of a bioelectronic patch having the conjugated polymer, polyaniline (PANI), as its conductive material.

EO, autoclave and γ-irradiation are examples of commonly used sterilization procedures in the clinic.^14,15^ The choice of a suitable sterilization technique is dictated by the medical implant and its constituent material. For a polymeric implant, careful consideration of the chemical structure of the polymer backbone should be made as the applied sterilization technique can react with the polymer. For instance, polyurethane that contains ester and urethane bonds in the backbone degrades significantly under γ-irradiation at a low dose of 25 kGy.^16^ EO is a chemical treatment shown to leave residues in the biomaterial, as well as affect the bioactivity and structural integrity of protein-based scaffolds.^15,17^ Similarly, the conjugated nature of the polymer in organic bioelectronics might be altered by the sterilization technique used, which may be detrimental to the device performance. Poly(3,4-ethylenedioxythiophene)-poly(styrene sulfonate) (PEDOT:PSS) electrodes sterilized by autoclave were shown to retain their electronic properties and surface morphology.^18^ However, when subjected to hydrogen peroxide gas plasma, the electronic and physical properties of the electrodes were significantly degraded. This was attributed to extensive crosslinking of the PEDOT:PSS chains, disrupting the conjugation length and thereby deteriorating the electronic and mechanical properties of the polymer.^18^ In contrast, autoclave sterilization of polypyrrole (PPy) electrodes resulted in complete loss of electrochemical activities and significant increase in impedance, whereas γ-irradiation at doses from 15 to 75 kGy did not affect the surface morphology, electrical, and electrochemical properties of the electrodes.^19^ Besides these two studies, the majority of the research that investigated *in vivo* implantation of conjugated polymers reported only the sterilization technique without any assessment of its effect on the polymer functional properties.^20–28^ It is clear from the brief literature that the small number of studies dedicated to understanding the relation between sterilization and device functional properties might be a contributing factor to the lack in advancement of organic bioelectronics in clinical applications.

Thus, the objective of this study was to investigate the effect of clinically approved sterilization techniques on the physical and electronic properties of organic bioelectronic devices as well as assess the inflammatory response caused by their *in vivo* implantation. We chose for our investigations a porous PANI-based bioelectronic patch previously developed by our group, driven by two of its distinctive properties: porosity and enhanced electronic stability in physiological environment.^29–31^ The porous nature of the patch is an engineering design that will aid in its *in vivo* integration. Porous biomaterials have been shown to integrate more efficiently with tissue by promoting cell diffusion, neovascularization, and thus leading to a reduced fibrotic capsule.^32^ Its demonstrated electronic stability in physiological conditions makes it the ideal bioelectronic interface for further testing *in vivo* and understanding the effect of sterilization technique on its physical and electronic properties. We fabricated the porous bioelectronic patch by synthesizing PANI doped with phytic acid on a porous chitosan film.^29^ We sterilized the patch by autoclave, EO and γ-irradiation of different doses. We investigated its mechanical and electronic properties before and after sterilization. We performed an *in vivo* study for 2 and 4 weeks to monitor the inflammatory response and determine the most effective sterilization technique for PANI.

## 2. EXPERIMENTAL SECTION

### 2.1. Materials

Chitosan (medium molecular weight, 85% degree of deacetylation), acetic acid, ammonium persulfate (APS), aniline, and phytic acid (50 wt% in water) were purchased from Sigma-Aldrich. Harris haematoxylin and eosin were purchased from Thermo Fisher Scientific, Hematoxylin solution Gill No. 3 and NovaRED™chromogen stain from Vector Laboratories, and Masson’s trichrome from Sigma Aldrich.

### 2.2. Fabrication of the porous PANI-based bioelectronic patch

The porous bioelectronic patch was fabricated following our previously published protocol ^30,31^, and as schematically presented in Figure 1A. Briefly, 1.6 mL of a stock solution of chitosan (1.6 wt% in 2 v/v% acetic acid solution) was drop-casted on a microscope slide (75 mm × 26 mm) and left to dry at room temperature for two weeks.^33–35^

**Figure 1.**
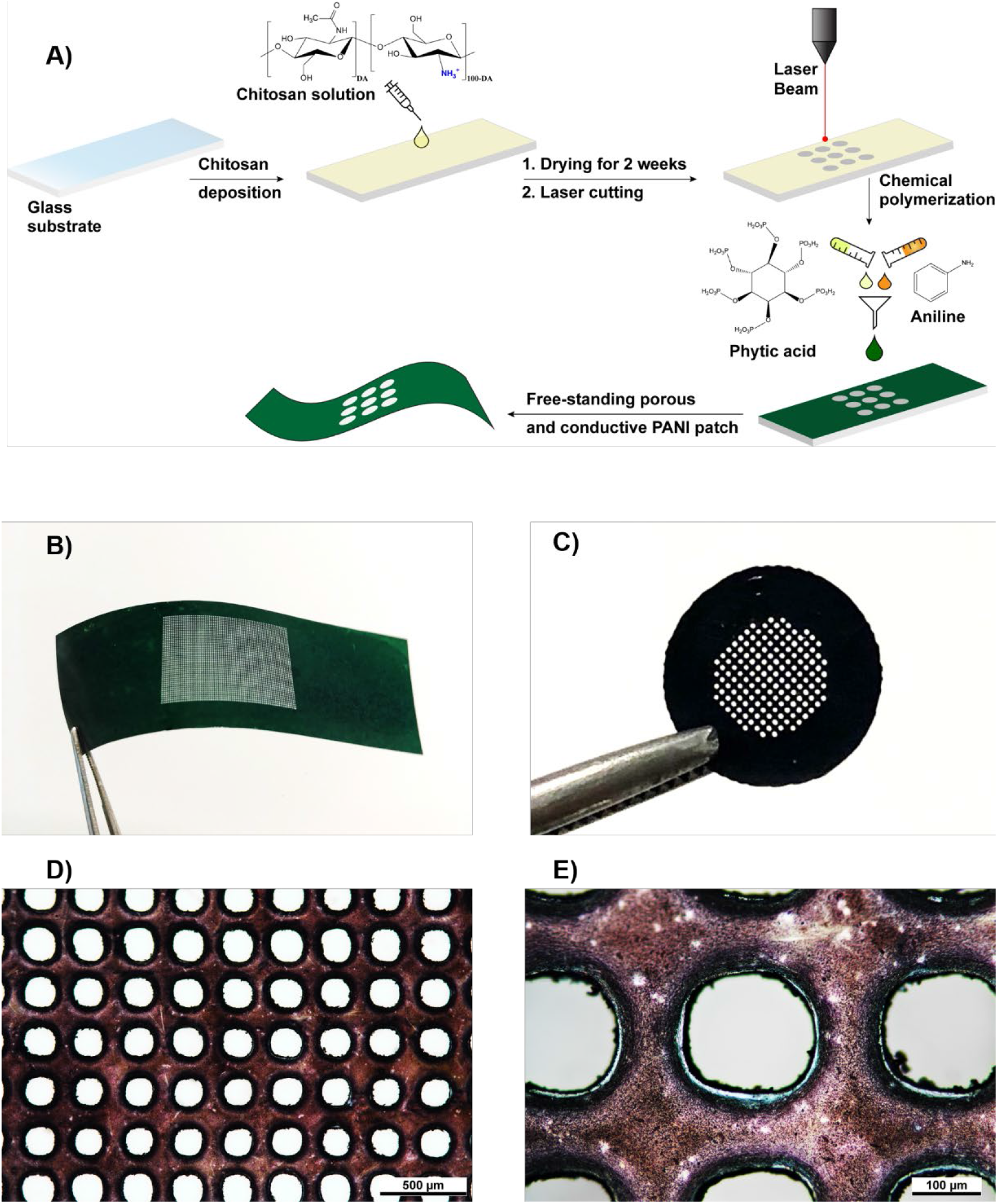
A) Schematic presentation of the fabrication process of the porous and conductive PANI patch. Photos of the free-standing PANI patch with rectangular (B) and round geometry (C). D and E) Optical microscope images of the pores created. Scale bars are 500 μm (D), and 100 μm (E).

A laser engraver (Trotec SP500, power 15%, speed 100%, and frequency 1 kHz) was used to create pores in the chitosan film. The porous films were then cut using the laser into two geometries: rectangular and circular. Rectangular films were 65 mm in length and 20 mm in width, with a central porous rectangular area of 20 mm in length and 14 mm in width (Figure 1B). Circular films had a diameter of 8 mm with a porous internal structure having a diameter of 4 mm (Figure 1C). The rectangular films were used for physical and electronic characterization, and the circular films for the *in vivo* experiments. Non-porous rectangular films (1 × 1 cm) were also prepared for X-ray photoelectron spectroscopy analysis and measurement of the sheet resistance. PANI was then polymerized by oxidative polymerization of aniline on the surface of the porous chitosan films.

In a typical procedure for the synthesis of PANI on the rectangular patches, ammonium persulfate (0.284 g) was dissolved in DI-H_2_O (1 mL). Phytic acid (0.92 mL) and aniline (0.46 mL) were mixed with DI-H_2_O (2 mL) in a separate vial. APS solution (0.80 mL) was then mixed with the aniline-phytic acid solution (2.27 mL) and evenly spread on the porous chitosan substrate. For the circular patches, these ratios were adjusted based on the surface area of the patch. The reaction was left for 3 hours to allow for the polymerization of aniline. After that, the patch was rinsed to remove any residual monomer and oxidant, as well as any polymer flakes that did not adhere to the chitosan film. The patch was soaked in deionized water (DI-H_2_O) overnight, dried at room temperature for one day, and stored without any further modification until used in the characterization experiments.

### 2.3. Optical and microscope images

The diameter of the pores and the distance between the pores were evaluated using a Nikon 600 optical microscope. Five porous PANI patches were used. Two different areas from each patch were selected and imaged. From each area, 5 pores were selected for measuring the diameter and distance using the Gwyddion software.

### 2.4. Sterilization

Three sterilization techniques were used: autoclave, EO, and γ-irradiation applied at 3 different doses. All patches were sterilized while still adhered on glass slides following the fabrication process.

#### 2.4.1. Autoclave sterilization

Patches, packaged in a double fold bag (Sterilope), were loaded in an autoclave chamber (Sanyo/Panasonic MLS-3871L autoclave) and were subject to a temperature of 121 °C and a pressure of 1.21 MPa for 10 min. The chamber was then cooled for 20 minutes before retrieval of the patches.

#### 2.4.2. EO sterilization

Patches were placed in petri dishes and exposed to ethylene oxide gas for 10 h, followed by 2 h degassing in the sterilizer chamber. Samples were then transferred to a vacuum oven at room temperature for an additional 24 h to remove residual EO.

#### 2.4.3. γ-irradiation

Patches were placed in petri dishes and stored in a locked zip bag for radiation exposure at a cobalt-60 γ-irradiation facility at the Australian Nuclear Science and Technology Organization. Each patch was placed in the irradiation room at pre-determined distances from the radiation source in order to irradiate samples at multiple doses simultaneously. Dosimetry was performed using calibrated poly(methyl methacrylate) (PMMA) dosimeters, and the average doses were found to be 15, 25, and 50 kGy.

### 2.5. Mechanical characterization

Tensile testing was performed on the rectangular patches (∼ 20 μm in thickness) before and after sterilization. The patches were wetted by DI-H_2_O to mimic physiological conditions before mounting them on steel grips of an Instron 5565 interfaced with Instron Bluehill software. The initial distance between the two grips was measured and a strain rate of 10 mm/min was applied. Ultimate tensile stress (UTS), Young’s modulus and elongation at break were estimated from the stress-strain curve. The Young’s modulus was calculated from the slope of the linear fit applied in the strain range between 0 to 5%. Five samples for each group were tested.

### 2.6. Electrical characterization

Sheet resistance (kΩ/□) of the patches was measured by a 4-probe station (Ossila Ltd.) The reported relative sheet resistance was calculated by dividing the sheet resistance of the sterilized patches by the sheet resistance of the non-treated patch.

### 2.7. X-ray photoelectron spectroscopy (XPS)

XPS was used to probe changes in the oxidation state and chemical structure of the patches subject to sterilization. XPS spectra were obtained using a Thermo Scientific ESCALAB250Xi spectrometer with a mono-chromated Al Kα x-ray source (energy 1486.68 eV), with a photoelectron take-off angle of 90 degrees. The survey spectrum for each sample was recorded using a pass energy of 100 eV in the range 0 to 1350 eV. This was followed by high resolution measurements with a pass energy of 20 eV for C 1*s*, N 1*s*, O 1*s*, S 2*p*, and P 2*p* core levels. The spectrometer was calibrated to Au 4*f*7 (83.96 eV), Ag 3*d*5 (368.21 eV) and Cu 2*p*3 (932.62 eV), and the binding energy reference was C 1*s* (284.6 eV) for adventitious hydrocarbon. Peaks were deconvoluted using the Avantage software 2.0 with the constraint of having the full widths at half maximum (FWHM) set at 1.5-1.7 eV for the N 1*s* peaks and 1.2-1.4 eV for the C 1*s* peaks, following values reported in the literature.^36–38^

### 2.8. In vivo study

#### 2.8.1. Animal procedure

The animal care and ethics committee at UNSW Sydney approved all animal work (ACEC:18/149B) on 8-week-old female C57/BL6 mice (Australian BioResources). Mice were placed under general anesthesia of isoflurane (1 L/min oxygen, 3-4% induction, 2-3% maintenance). The back of each mouse was shaved, and surgical site disinfected with betadine and isopropanol wipes thrice. Buprenorphine was administered at 0.1 mg/kg, subcutaneously preoperatively. One surgical skin incision was made below the neck and subcutaneous pockets, towards the left front leg, left hind leg, and right front leg, were opened with blunt forceps. One randomized circular patch was implanted in each subcutaneous pocket (n = 5/timepoint). The incisions were closed with surgical clips, and wound adhesive was applied at the wound site. Surgical clips were removed at day 10 post-surgery. Postoperatively, animals were housed with unlimited access to water and food.

#### 2.8.2. Histology analysis

Patches together with the surrounding tissues were explanted after 2 and 4 weeks and were fixed in 10% neutral-buffered formalin for 24 hours, dehydrated in 70% ethanol and embedded in paraffin wax. Thereafter, the sections were stained with haematoxylin and eosin (H&E), Masson’s trichome as previously described.^39^ Briefly, following rehydration, slides were stained with Harris haematoxylin for 5 min, followed by eosin for 4 min, before dehydration and mounting. Masson’s trichrome staining was performed following manufacturer’s standard procedure. Briefly, samples were mordanted in preheated Bouin’s solution at 56 °C for 1 hr and washed with tap water. The samples were then stained with Hematoxylin solution Gill No. 3 for 5 min, Biebrich scarlet-acid fucshin for 5 min, phosphotungstic/phosphomolybdic acid solution for 5 min, and lastly aniline blue solution for 5 min with 5 min tap water washes in between each stain. The samples were placed in 1% acetic acid for 1 min before dehydration and mounting.

For immunohistochemical analyses, antigen epitope retrieval was performed by immersing the slides, following rehydration, in 0.01 M sodium citrate (pH 6), followed by either heat treatment in a decloaking chamber (Applied Medical) at 120 °C for 4 min or by incubation with 0.05% wt/v proteinase XXIV in Tris-buffered saline (TBS, 0.05 M Tris, 0.15 M NaCl pH 7.6) at 37 °C for 15 min. The slides were then rinsed with TBS followed by blocking with 10% v/v goat serum in TBS for 1 h at room temperature. The slides were incubated with primary antibody diluted in 1% v/v goat serum in TBS at 4 °C for 16 h. Slides were then washed twice with TBST (TBS with 0.05 % wt/v Tween 20) before incubating with the appropriate biotinylated secondary antibodies (GE Healthcare, 1:500) for 1 h at room temperature. The slides were washed twice with TBST then incubated for 30 min with streptavidin-HRP (1:250), rinsed four times with TBST before color development with NovaRED™chromogen stain. The slides were then counterstained with Hematoxylin solution Gill No. 3 for 10 s and rinsed with DI-H_2_O. All samples were imaged using Aperio ScanScope XT scanner.

### 2.9. Statistics

Statistical analysis was performed using one-way ANOVA with Tukey’s multiple comparison test (Graphpad Prism). A *p* value < 0.05 was considered statistically significant.

## 3. RESULTS AND DISCUSSION

### 3.1. Porous PANI-based bioelectronic patch

We introduced porosity in the bioelectronic patch using a laser engraver technology since porosity is a well-established design parameter for implantable devices shown to reduce the inflammatory response.^40^ As shown in Figures 1D and 1E, the laser engraver successfully created regular and circular pores in the patch, achieved by melting the chitosan polymer. The pore shape was maintained following the polymerization of aniline on the surface of the chitosan substrate. The final average pore diameter was estimated at 169.6 ± 6.9 μm with an average distance of 101.8 ± 11.0 μm between the pores, and a pore density of ∼ 28%. The porous patches retained their flexibility and mechanical integrity. The optimal size and density of pores for the ideal scaffold depend on type of the host tissue.^41^ The laser engraver technology we used enables the design of porous patches where the size of pores and distance between pores can be controlled independently, as we have previously shown.^31^

## 3.2. Mechanical properties

We assessed the impact of sterilization on the mechanical properties of the patches using tensile testing. Autoclave sterilization resulted in numerous cracks in the patch. The patches are adhered to the glass substrate following fabrication. We speculate that subjecting the patches to high temperatures (121 °C) might have resulted in the rapid evaporation of residual water from the films. Having the patch adhered to the glass slide might have caused restriction on the mobility of the polymer chains.^42^ and thus caused significant changes in its internal morphology and shrinkage in response to the fast evaporation rate. Consequently, no further characterization was performed on patches sterilized by this technique.

Figure 2 presents the mechanical properties of the non-treated, EO, and γ-irradiated patches.

**Figure 2.**
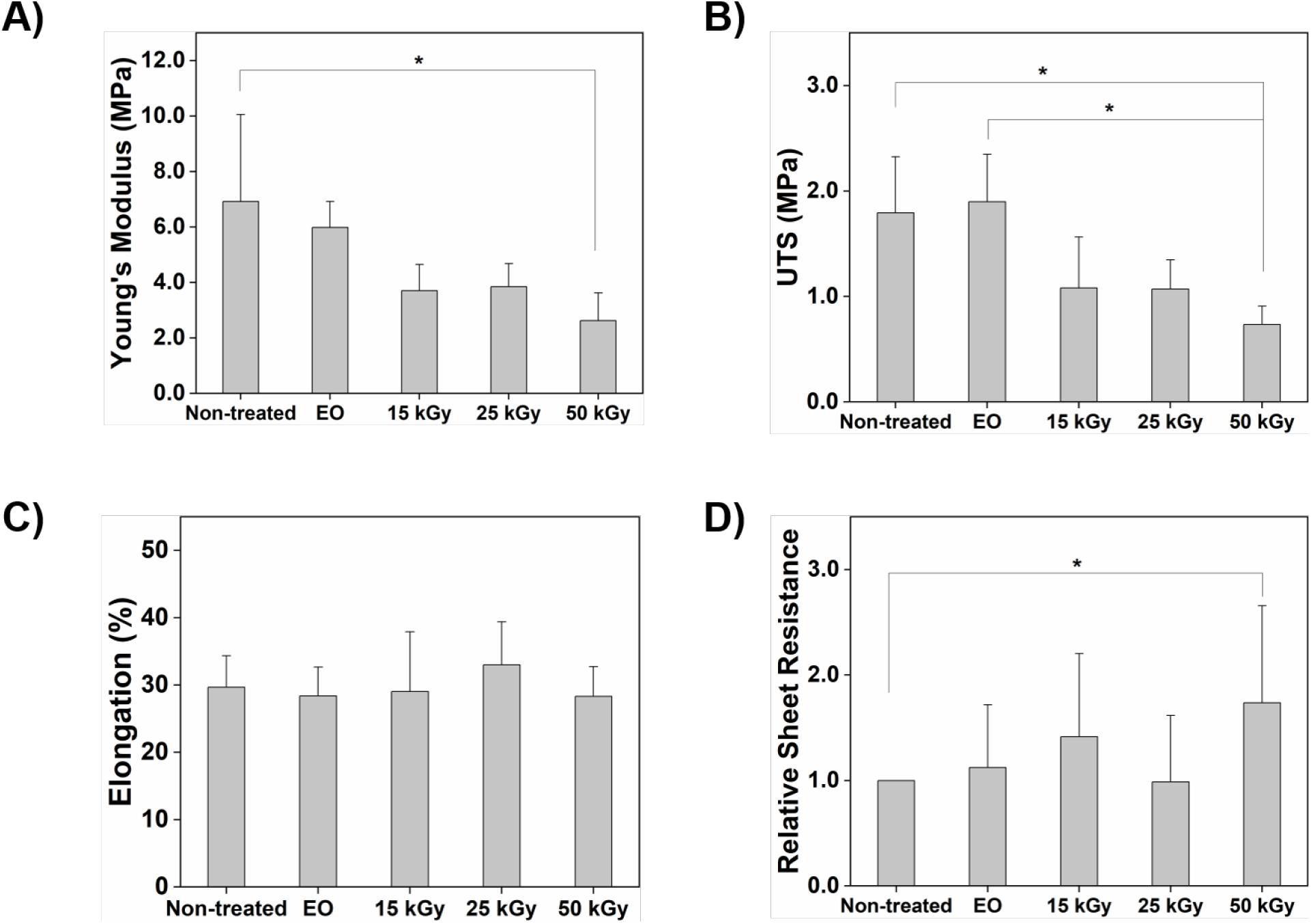
Physical and electronic properties of the non-treated and sterilized patches: A) Young’s modulus, B) ultimate tensile stress (UTS), C) elongation at break, and D) Relative sheet resistance. (*) denotes a significant difference. Error bars represent the standard deviation for n= 5.

Non-treated and EO patches exhibited similar values of Young’s modulus, 6.91 ± 3.14 and 5.98 ± 0.94 MPa respectively. The Young’s modulus for the γ-irradiated patches varied from 3.70 ± 0.94 MPa for patches treated with 15 kGy dose to 3.85 ± 0.83 MPa, and 2.62 ± 1.00 MPa, for patches treated with 25 kGy and 50 kGy doses, respectively (Figure 2A). While the results suggest a decreasing trend in the Young’s modulus, a significant difference was only detected in the Young’s modulus of patches irradiated with 50 kGy compared to that of the non-treated patches (Tukey’s post-test analysis, *p* < 0.016). A similar trend was observed for the ultimate tensile stress (Figure 2B), with a significant drop observed only for the ultimate tensile strength of the patches treated with 50 kGy in comparison to non-treated (Tukey’s post-test analysis, *p* < 0.0117) and EO patches (Tukey’s post-test analysis, *p* < 0.0102). Elongation at break was similar for all patches.

Changes in the mechanical properties of polymers subject to γ-irradiation have been attributed to two competing mechanisms: chain scission and crosslinking of the polymer chains.^43,44^ Chain scission will result in a reduction in the mechanical properties such as the Young’s modulus and ultimate tensile stress, whereas crosslinking is expected to improve the mechanical properties. Our results indicate that the 50 kGy dose induced significant drop in the mechanical properties of the films. This suggests that chain scission is most likely the predominant mechanism at this high dose of irradiation, although non-uniform crosslinking can not be ruled out. γ-irradiation at 40 kGy dose has been shown to decrease the ultimate tensile strength of chitosan films, attributed to scission of the 1-4 glycosidic bond, and inhomogeneous crosslinking of the polymer chains that create non-uniform stress distribution in the film.^45–47^ Similarly, chain scission has been reported in γ-irradiated polyaniline at doses as low as 10 kGy, producing free radicals that can recombine to form a crosslink or react with oxygen when the films are exposed to air.^48^ Here, our patches could sustain γ-irradiation up to 25 kGy without any significant changes in their mechanical properties. The PANI is polymerized on the surface of the chitosan film and it appears that the mechanical properties are governed by the chitosan polymer rather than PANI.

### 3.3. Electrical characterization

We tested the impact of sterilization on the electronic properties of the patch by measuring sheet resistance using a 4-probe setup. Figure 3 presents the relative sheet resistance of the sterilized patches in comparison to the non-treated patches. Patches sterilized with EO and γ-irradiation at 15, and 25 kGy doses exhibit a sheet resistance comparable to the non-treated patch, indicating that EO sterilization and γ-irradiation at these doses have no effect on the electronic properties of the patch. In contrast, the sheet resistance of the 50 kGy γ-irradiated patches increases by ∼74% relative to the original value of the non-treated patches (Tukey’s post-test analysis, *p* < 0.0476). This result is in agreement with the tensile testing that revealed a reduction in the mechanical properties of the patches irradiated with 50 kGy dose, and confirms that chain scission, and/or non-uniform crosslinking, of polyaniline might be occurring when irradiated with the 50 kGy dose. Chain scission and crosslinking will cause the conjugation length of the polymer to be reduced, hindering intra- and interchain transport and thus accounting for the drop in electrical properties.^49^

**Figure 3.**
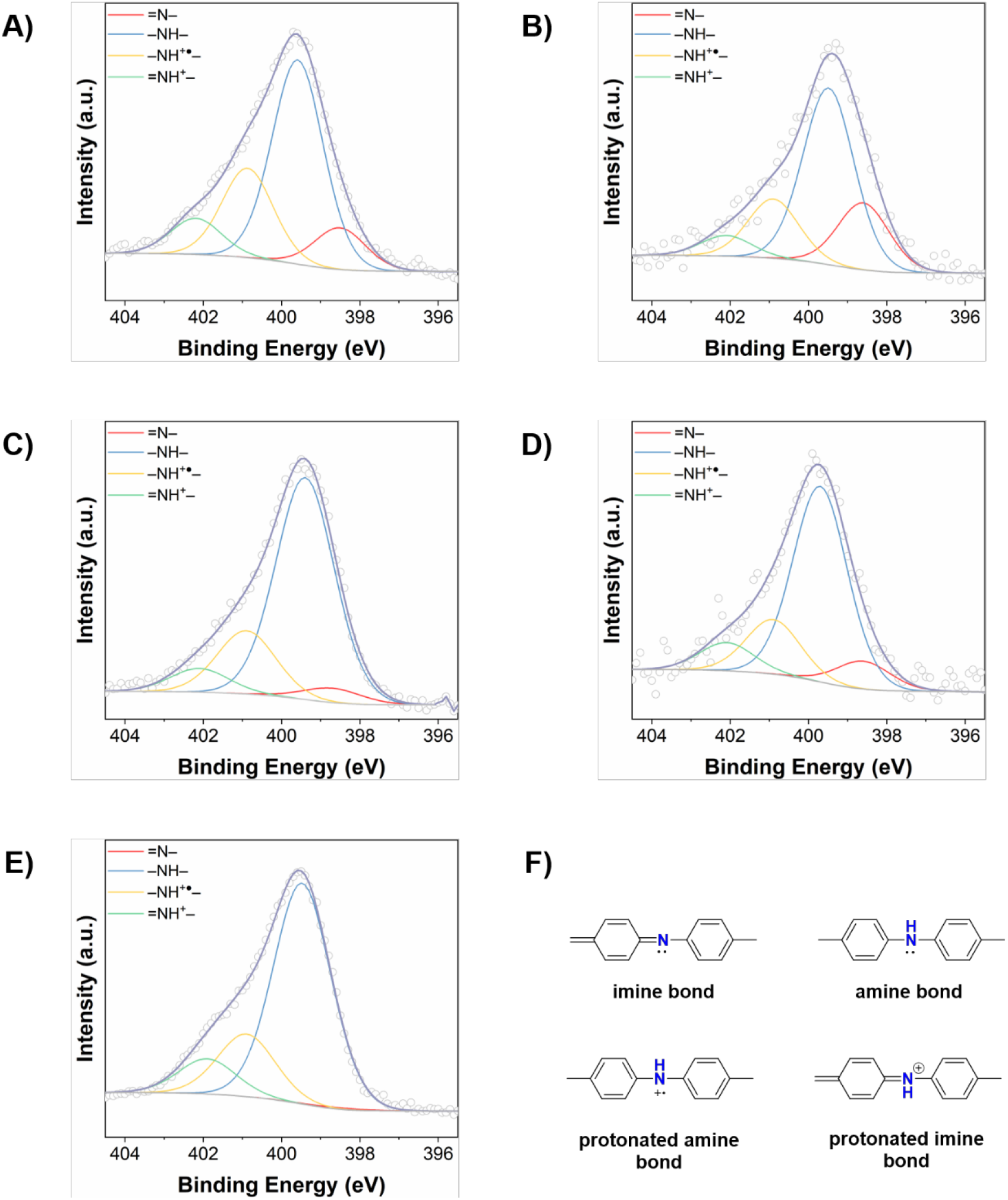
The N 1*s* deconvoluted spectra of A) non-treated, B) EO sterilised, C) 15 kGy γ-irradiated, D) 25 kGy γ-irradiated, and E) 50 kGy γ-irradiated patches. F) Schematic presentation of the type of nitrogen bonds present in polyaniline.

Decrease in electric properties in response to γ-irradiation at doses ranging from 5 to 10^3^ kGy have been reported for drop-casted polyaniline on a polyester substrate^48^ and spin coated polyaniline films.^50^ Of note, the electric properties were shown to deteriorate if γ-irradiation occurs in air due to oxidation of the film surface ^48^. However, other studies have reported an enhancement in the electric properties of γ-irradiated polyaniline, in a pellet form or as a composite, even at doses as high as 400 kGy.^51,52^ Proposed mechanism for the improvement in conductivity is attributed to an increase in radical concentration. Various parameters, such as the exposure time to the radiation and the thickness of the polymer, should be considered before comparison can be made between different studies. To date, the literature concerning the impact of irradiation on conducting polymers remains limited.

### 3.4. XPS analysis

We used XPS analysis to probe changes in the chemical structure of PANI following sterilization. There are large variations in the assignment of the resolved peaks of the C 1*s* and N 1*s* of polyaniline in the literature.^36–38,53–56^ Here, the positions of our fitted peaks and FWHM values correspond closely to those reported by Kumar et. al. ^36,37^ Additionally, our patch contains chitosan and phytic acid with overlapping elements with polyaniline: nitrogen and protonated NH_3_^+^, carbon, and oxygen. Thus, due to the complexity of the chemical composition of the patch, we limit our discussion of the XPS results to a qualitative assessment of the assigned peaks, rather than an absolute quantification of their intensities. However, we do present the relative intensities in Tables S1 and S2 to provide a better insight of the data. We also include chemical structures to present the different groups of nitrogen and carbon that we have resolved in the XPS spectra (Figures 3F, Figure S1 and Table S3).^37^

The N 1*s* core spectra and its relative fitting are shown in Figure 3. The nitrogen in a doped PANI polymer typically has four chemical environments: imine (=N–) at ∼398.5 eV, amine (–NH–) at ∼399.5 eV, protonated amine (–NH^+●^–) at ∼401.0 eV, and protonated imine (=NH^+^–) at ∼402.0 eV. The protonated species are attributed to polarons and bipolarons, respectively, in the polymer backbone ^37,57^. Inspection of the figures reveals that the deconvoluted N 1*s* envelopes are similar for all patches, non-treated and sterilized (Figure 3), except for the 50 kGy patch that had a negligible peak corresponding to the imine (∼ 0.47%) (Figure 3E).

The patches had a slight variation in their doping ratio (∼ 27 to 33%, Table S1); nevertheless, this change had no effect on the electronic properties of the patches as shown by the relative sheet resistance results. Again, the 50 kGy was the exception, exhibiting an increase in its resistance while still having a doping ratio of ∼ 30% comparable to the other patches.

We fitted the C 1*s* core level spectra presented in Figure 4 to further understand any changes that might have occurred in the patches, in particular the 50 kGy patch. We first consider the C 1*s* spectrum of the non-treated patch that was deconvoluted into five peaks (Figure 4A), similar to analysis performed by Kumar and co-authors ^36,37^. The first four peaks with FWHM of 1.4 eV correspond to: 1) C–C and C–H of the aromatic groups (∼ 284.7 eV), which form the backbone of the polymer and have the highest relative intensities, 2) C–N and C=N (∼ 285.7 eV) assigned to the carbon bonded to the neutral nitrogen atoms (amine and imine), 3) C–N^+●^ (286.7 eV) assigned to the carbon bonded to the positively charged amine, and 4) C–N^+^ (∼ 287.7 eV) assigned to the carbon bonded to the protonated imine (Figure S1). The peak with the highest binding energy at ∼ 288.8 eV and FWHM of 1.7 eV is attributed to C=O bonds that can form during polymerization or following exposure of the patch to air ^53,58^. The C 1*s* spectra of the sterilized patches could only be fitted to four peaks if we wanted to maintain the FWHM constant between all the samples (Figures 4B-E). The peaks were assigned as follows: =CH– at ∼ 284.7 eV, C–N, C=N, and C–N^+●^ at ∼ 286.1 eV, C–N^+^ at 287.1 eV, and C=O at ∼ 288.8 eV. Monkman and co-authors ^38^ reported that the C–N, C=N, and C–N^+●^ can be at very close binding energy based on bond potential calculations and thus might result in an unresolved peak ∼ 286 eV. They did their analysis on electropolymerized polyaniline films, similar to films studied by Kumar et. al. Based on these two studies, it is not clear why the deconvolution of C 1*s* can vary between 3 or 4 peaks despite that both analyses were performed on electropolymerized polyaniline. Thus, we conclude that the difference in the peak fitting between the non-treated patch and the sterilized patches is not related to the sterilization processes applied.

**Figure 4.**
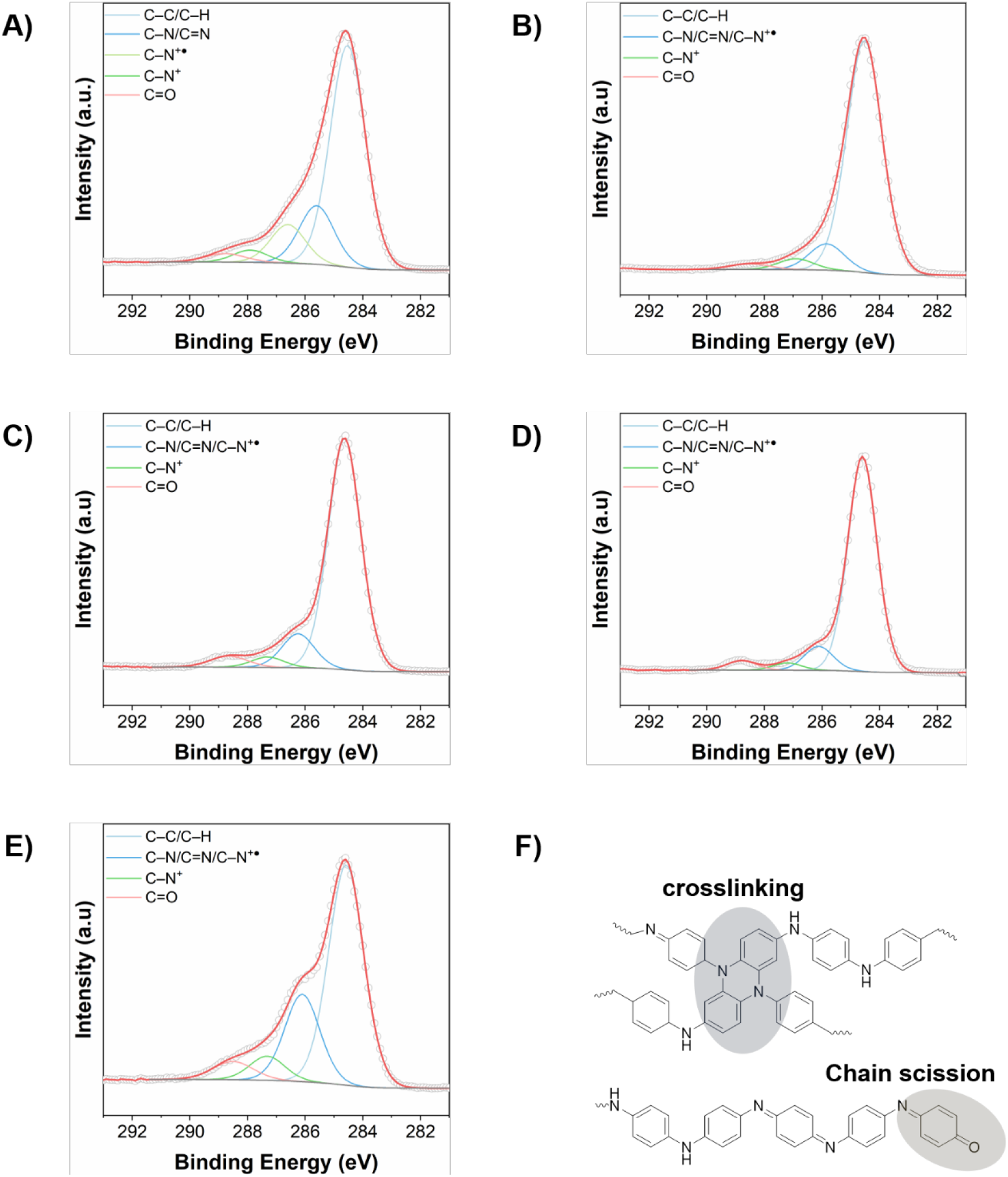
The C 1*s* deconvoluted spectra of A) non-treated, B) EO sterilised, C) 15 kGy γ-irradiated, D) 25 kGy γ-irradiated, E) 50 kGy γ-irradiated patches. F) Schematic presentation of potential crosslinking and chain scission that might have occurred in the 50 kGy irradiated patch. For simplicity, polyaniline is presented in its neutral state.

One distinctive difference between the patches is the intensity of the carbon bond in the ring and the intensity of the carbon bonded to neutral nitrogen atoms (Table S2 and Figure S1). For the non-treated patch (Figure 4A) and the patch irradiated with 50 kGy (Figure 4B), the intensity of the carbon bonded to nitrogen is higher than expected for polyaniline in which the ratio of C/N is 6. Our values are similar to those reported in the literature where this increased intensity has been attributed to hydrocarbon impurities.^36,37^ The non-treated patch was stored under non-sterile conditions and was exposed to the environment, which might explain the deposition of impurities on the surface. The sterilized patches were packaged immediately after fabrication and remained in sealed containers after being sterilized until the XPS measurements were performed. This might have lowered the impurities on the surface as reflected by the lower intensity of the peak corresponding to carbon bonded to nitrogen atoms for the EO, 15, and 25 kGy patches. We speculate that besides impurities, the increased intensity in the 50 kGy can be attributed to the formation of new C-N bonds assigned to tertiary amines (Figure 4F). As discussed earlier, the 50 kGy dose might be inducing chain scission and non-uniform crosslinking. The radicals that form upon chain scission can recombine to form new bonds such as tertiary amines resulting in crosslinking or can react with oxygen to form new carbonyl bonds (Figure 4F). Tertiary amine formation on the expense of imine bonds have been reported in aged polyaniline, with the binding energy of the tertiary amine bonds overlapping with the protonated amine.^54,55,59^ Indeed, the imine peak almost disappeared in the N 1*s* spectrum of the 50 kGy patch (Figure 3E), confirming that tertiary amine and thus crosslinking might have occurred when irradiated at this dose. Additionally, the intensity of the carbonyl peak is highest for the 50 kGy patch. Taken together, XPS analysis reveals that the 50 kGy dose is the only sterilization treatment that induces changes in the chemical structure of the polyaniline. However, it should be noted that XPS is a surface analysis and the bulk of the material might not reflect these changes.

Overall, the N 1*s* spectra indicates that the sterilization procedures do not result in full deprotonation of the PANI polymer and thus its conversion to a reduced or neutral state. This is in agreement with a previous study that showed that a reduction in conductivity of γ-irradiated polyaniline is attributed to chain scission and not due to de-protonation of the polymer, as demonstrated using Raman and UV-vis spectroscopy.^48^ The C 1*s* spectra points towards the formation of new bonds in the patches irradiated with 50 kGy. Overall, the XPS data appears to be in agreement with the mechanical and sheet resistance data. The protonated nature of all the patches indicates that the patch remains doped and thus conductive. Changes observed in the N 1*s* and C 1*s* spectra of the 50 kGy patch are in agreement with the reduced mechanical properties and increased sheet resistance.

### 3.5. In vivo studies

Our characterization studies revealed that the 50 kGy dose had detrimental effect on the physical and electronic properties of the patch; thus, we eliminated this dose from the *in vivo* study. We implanted the sterilized patches in subcutaneous pockets in mice to assess the impact of different sterilization techniques on *in vivo* performance of the patches, including surgical handling, inflammatory responses, and tissue integration at 2- and 4-weeks post-implantation. At the time of implantation, we observed that the EO-sterilized patches were flexible, easy to handle, and position in the subcutaneous pockets. In contrast, the γ-irradiated patches were difficult to handle and position precisely in the subcutaneous pockets. The patches were handled well *in vivo*, with all animals surviving the surgical procedure and subsequent incubation period, and no obvious tissue irritation or inflammation observed at the surgical site macroscopically. At weeks 2 or 4 post-implantation, we explanted the patches along with the surrounding tissue and processed for histological and immunohistochemical analyses. Representative images of H&E stained EO, 15 kGy and 25 kGy γ-irradiation treated samples are presented in Figure 5A.

**Figure 5.**
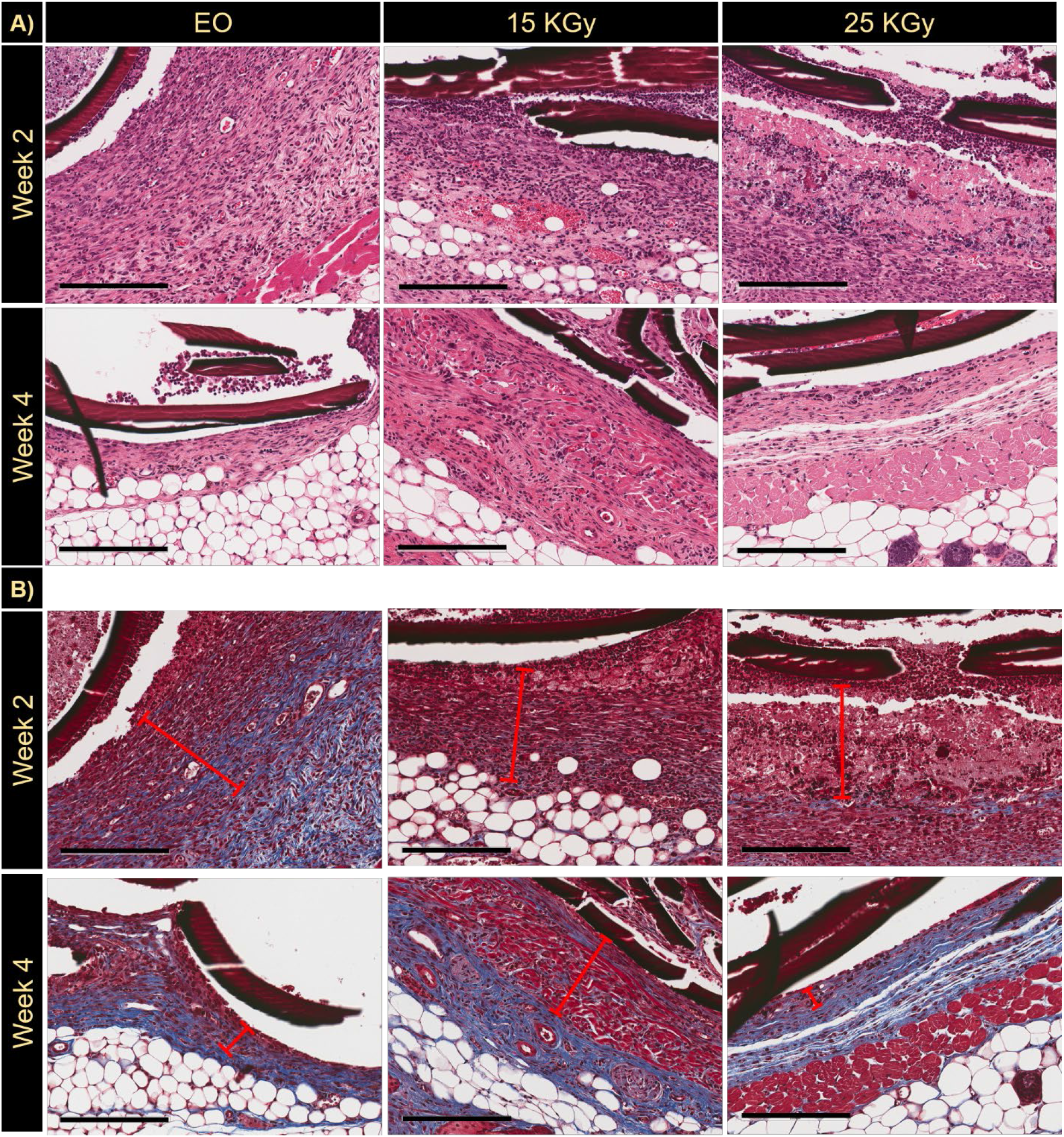
A) Hematoxylin and eosin (H&E) and B) Masson trichome staining after 2- and 4-week implantation of PANI patches treated with EO sterilization, 15 kGy and 25 kGy γ–irradiation. Scale bars are 200 μm.

The patches were integrated with the mouse skin during explantation and histologically appeared as dense pink films (Figure 5). The films did not appear as one continuous layer in histological section due to their porous structure. The tissue did not seem to be intimately associated with the patches throughout and this is believed to be an artefact of the histological processing due to the mechanical mismatch between the patches and the surrounding tissue, causing tissue to pull away from the films during sectioning. We observe at week 2 an obvious cellular immune response surrounding all the patches, seen as tissue densely populated by small cells with round nuclei. By week 4 this response was drastically reduced, with the tissue surrounding the implants becoming less cellular with more elongated cell nuclei typically associated with fibroblastic cells.

To probe the composition of the tissue surrounding the implants, tissue sections were stained with Masson trichome where collagen deposition appears blue as shown Figure 5B. The cellular response to implants was observed as dark pink stained region in proximity of the implant and is marked with red lines in the figures and is observed to reduce in thickness and change in composition by week 4 from a highly cellular tissue to collagen-rich tissue. We confirm this observation by quantifying the thickness of this region as presented in Figure 7A. The thickness of the immune/fibrotic response revealed that the type of sterilization technique did not influence the acute immune response, with all patches surrounded with a cellular tissue of ∼200 μm at week 2. However, the period over which the implant remained in the mouse had a significant impact on the capsule thickness, decreasing significantly for all patches after week 4 (Tukey’s post-test analysis, *p* < 0.0001). Similarly, the sterilization method and the dose appear not to impact the capsule thickness at week 4 as indicated by the Tukey’s post-test analysis (*p* ≥ 0.9471)

We performed immunostaining for CD31 to probe the vascularization of the tissue surrounding the implant. CD31 staining reveals that the tissues are well vascularized as presented in Figure 6.

**Figure 6.**
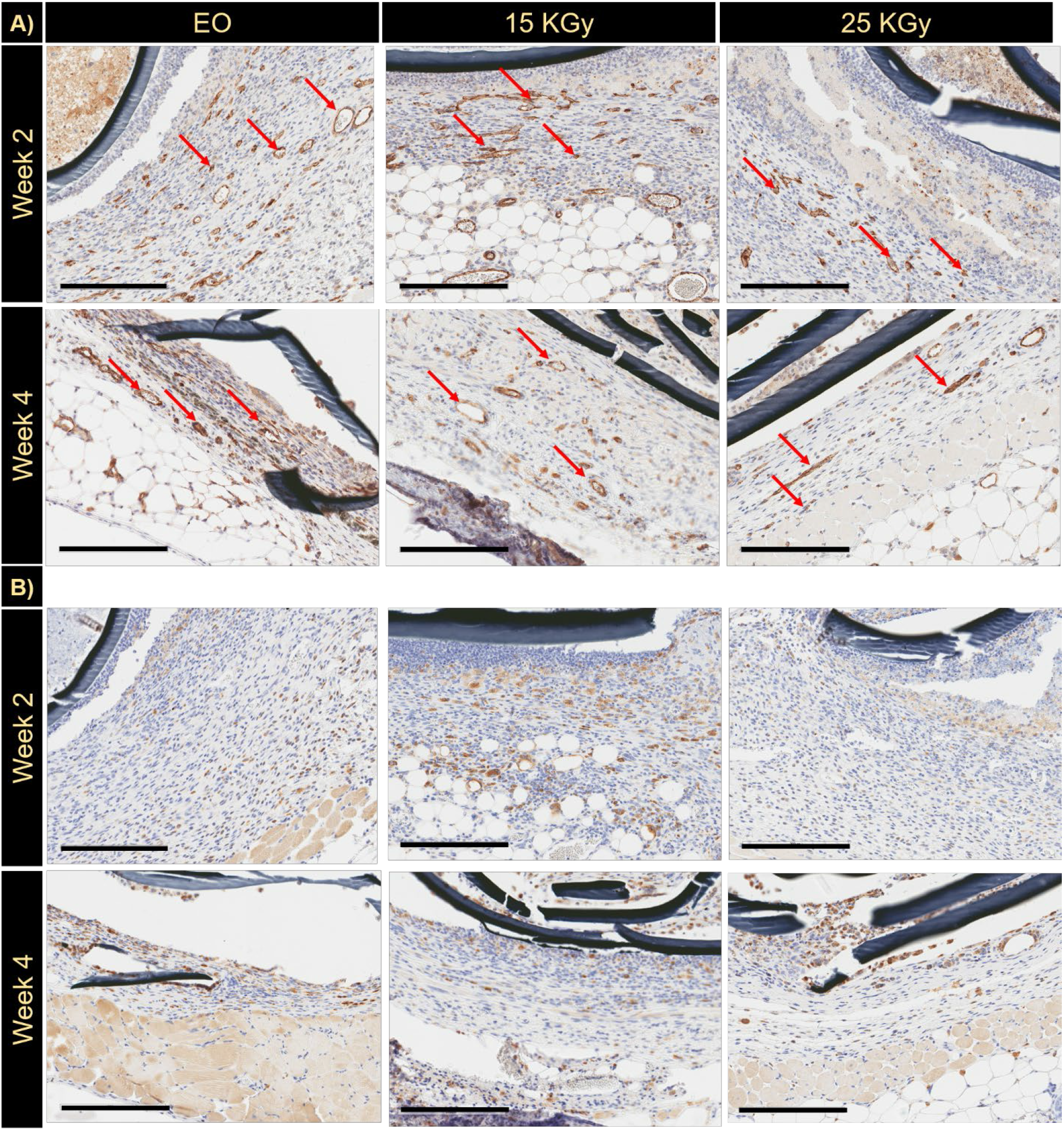
A) CD31 and B) CD68 stained samples after 2- and 4-week implantation of PANI patches treated with EO sterilization, 15 kGy and 25 kGy γ–irradiation. The red marker highlights the inflamed area. Scale bars are 200 μm.

**Figure 7.**
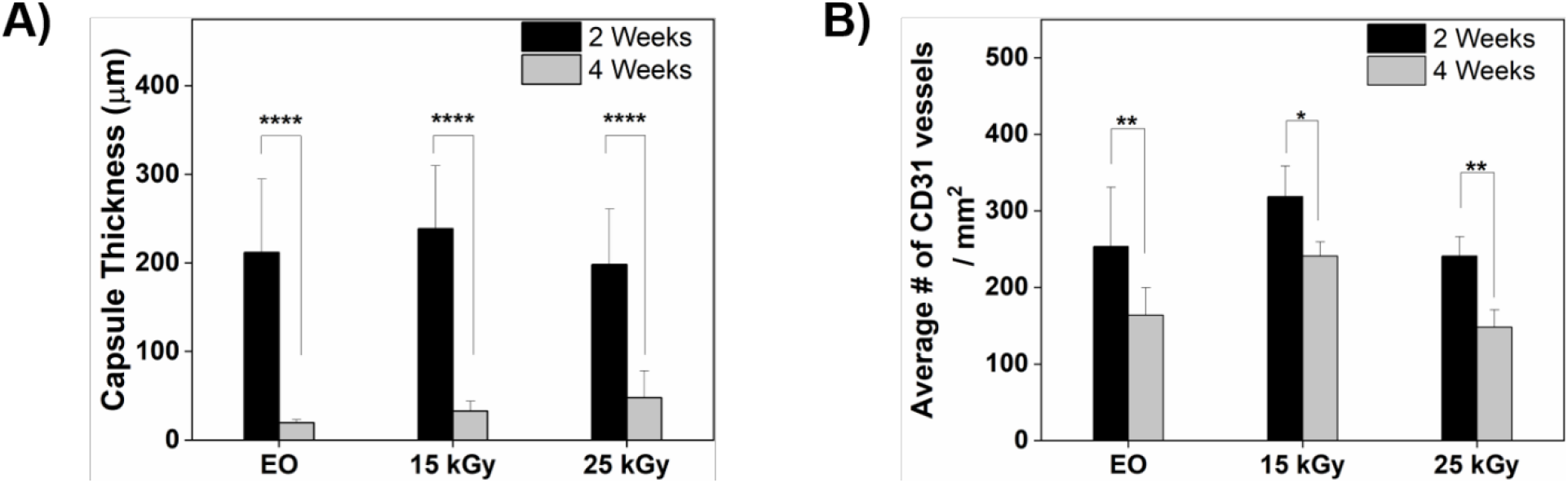
The thickness of the fibrotic capsule (A) and the average # of CD31 vessels (B) estimated at 2 and 4 weeks.

Vessels, highlighted with red arrows in the figures, are identified by the typical round shape that the staining technique marks with a characteristic brown color. The vessel density appeared significantly reduced by week 4 (Tukey’s post-test analysis, *p* < 0.0253) consistent with tissue maturation from a highly cellular response to extracellular matrix deposition (Figure 7B). No abnormalities were identified in proximity of the implant, especially close to the PANI patch. Immunostaining with CD68 revealed a moderate macrophage response that reduced by week 4, in agreement with the H&E and Masson trichome observations.

The patch but in its non-porous form has been previously implanted in a cardiac model following disinfection with ethanol.^29^ After two weeks implantation, the non-porous patch was surrounded by a fibrotic capsule which was 830 μm in thickness.^29^ The immune response observed and the use of ethanol in this early study promoted us to develop the next iteration of the PANI patch with engineered porosity and sterilized using clinically acceptable techniques (EO or γ-irradiation), unlike previously ethanol. In this study, the two time points were chosen in order to investigate both acute and chronic inflammatory response, respectively.^60,61^ The cellular/fibrous response surrounding the implants was ∼ 200 μm thick at 2 weeks and ∼ 30 μm thick at 4 weeks, suggesting an improved response relative to the previous study. While it is difficult to compare biological responses between models, these observations suggest the porous PANI patch is promising in terms of tissue response and fibrous encapsulation.

## 4. CONCLUSIONS

We have investigated the impact of sterilization techniques on the physical and electronic properties of a porous conductive PANI patch. Autoclave sterilization resulted in significant cracking in the patch and thus was eliminated from the study. EO and γ-irradiation at 15 and 25 kGy doses did not impact the mechanical and electronic properties of the patch. In contrast, γ-irradiation at 50 kGy dose resulted in a significant drop in the Young’s modulus and ultimate tensile stress, and a significant increase in its sheet resistance. Similarly, XPS analysis revealed significant changes in the N 1*s* and C 1*s* spectrum only for patches irradiated at 50 kGy dose. We speculate that the 50 kGy dose might be causing chain scission and non-uniform crosslinking, which in turn resulted in deterioration in the mechanical and electronic properties of the patch. A low inflammatory response was observed for the EO, and 15 and 25 kGy γ-irradiated patches, with the fibrotic capsule significantly receding after 4 weeks implantation for all three sterilization procedures. Based on these results, the 50 kGy dose is deemed detrimental to the patch properties, and thus not suitable for its sterilization. However, the EO, 15 and 25 kGy appear to be suitable for the sterilization of the patch, making it a clinically viable bioelectronic device.

## ASSOCIATED CONTENT

### Supporting Information

The following files are available free of charge.

Table S1: Peak intensities of the deconvoluted N 1s peaks.

Table S2: Peak intensities of the deconvoluted C 1*s* peaks.

Table S3: Binding energies (eV) corresponding to the bonds formed by the nitrogen and carbon elements as assigned in peak deconvolution of the XPS data.

Figure S1: Chemical structures of PANI highlighting the type of bonds that nitrogen and carbon form in the backbone.

## AUTHOR INFORMATION

### Author Contributions

The manuscript was written through contributions of all authors. All authors have given approval to the final version of the manuscript.

## ACKNOWLEDGMENT

DM, DO, and AL would like to acknowledge the Australian Research Council, Discovery Project Grant DP190102560, for funding this research. JR-K would like to acknowledge funding support from the Australian Research Council (DP150104242) and the NSW Health Cardiovascular Investigator Development Grant. JR-K was partially supported by the Heart Foundation of Australia Future Leader Fellowship (101896). KL was supported by the Australian Government Research Training Program Scholarship. Histological staining and imaging were performed at the Biomedical Imaging Facility (BMIF), and XPS was performed at the Surface Analysis laboratory. Both laboratories are part of the Mark Wainwright Analytical Center at UNSW Sydney and in part-funded by the Research Infrastructure program at UNSW. The authors would like to thank Ms Naomi Craig for technical assistance with the animal study, and Mr. Tim Cole from the Sterilizing Services Department at Prince of Wales Hospital, Randwick NSW for performing the EO sterilization.

## ABBREVIATIONS

EO: ethylene oxide
PANI: polyaniline
UTS: ultimate tensile stress
XPS: X-ray photoelectron spectroscopy

