## Supplementary Information for "Impact of sterilization on a conjugated polymer based bioelectronic patch"

**Table S1:** Peak intensities of the deconvoluted N 1*s* peaks.

| Patch | C=N– | C–NH– | C–NH•+– | C=NH+ | % doping (sum of protonated species) |
|---|---|---|---|---|---|
| **PANI** | 11.23 | 54.97 | 24.06 | 9.75 | 33.81 |
| **EO** | 9.09 | 63.87 | 17.84 | 9.20 | 27.04 |
| **15 kGy** | 4.47 | 68.63 | 19.57 | 7.33 | 26.90 |
| **25 kGy** | 9.09 | 63.87 | 17.84 | 9.20 | 27.04 |
| **50 kGy** | 0.47 | 69.01 | 19.59 | 10.97 | 30.52 |

**Table S2:** Peak intensities of the deconvoluted C 1*s* peaks.

| Patch | C=C | C–NH– | C=N– | C–NH•+– | C=NH+ | C=O |
|---|---|---|---|---|---|---|
| **PANI** | 64.96 | 17.19 | | 11.30 | 3.61 | 2.93 |
| **EO** | 83.76 | | 9.43 | | 2.90 | 3.90 |
| **15 kGy** | 77.92 | | 12.09 | | 5.96 | 4.03 |
| **25 kGy** | 83.76 | | 9.43 | | 2.90 | 3.90 |
| **50 kGy** | 62.41 | | 24.73 | | 6.86 | 5.60 |

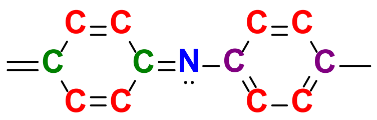

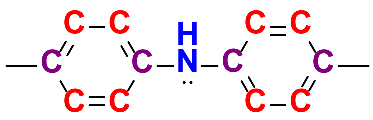

**(1)** **(2)**

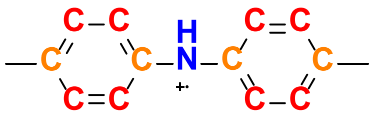

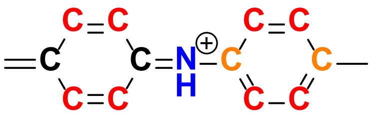

**(3)** **(4)**

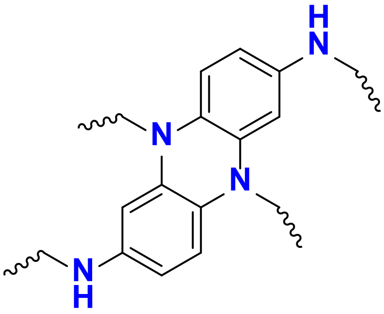

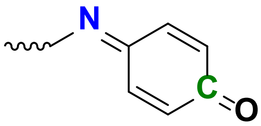

**(5)** **(6)**

**Figure S1**: Chemical structures of PANI highlighting the type of bonds that nitrogen and carbon form in the backbone.

**Table S3:** Binding energies (eV) corresponding to the bonds formed by the nitrogen and carbon elements as assigned in peak deconvolution of the XPS data.

| | | Binding energy (eV) corresponding to the nitrogen element[1–6] | Binding energy (eV) corresponding to the carbon element[1–4] |
|---|---|---|---|
| (1) | C=N— | ~ 398.5 | ~ 285.7 for PANI<br>~ 286.1 for EO sterilised and γ-irradiated patches |
| (2) | C—NH— | ~ 399.7 | ~ 285.7 for PANI<br>~ 286.1 for EO sterilised and γ-irradiated patches |
| (3) | C—NH•+— | ~ 400.9 | ~ 286.8 for PANI<br>Unresolved peak for EO sterilised and γ-irradiated patches[4] |
| (4) | C=NH⊕ | ~ 402.0 | ~ 287.1 |
| | C= | | ~ 284.7 |
| (5) | N | ~ 400.9 (overlap with the protonated amine)[7,8] | |
| (6) | C=O | | ~ 288.8 |
